# A chromosome-scale genome assembly for Clark’s Nutcracker (*Nucifraga columbiana*)

**DOI:** 10.1101/2025.11.06.687043

**Authors:** Peter A. Innes, Matt Carling, Ethan Linck

## Abstract

Focal and intelligent passerine songbirds, corvids (Aves: Corvidae) have served as models for research on the genomics of hybridization and cognition. Clark’s Nutcracker (*Nucifraga columbiana*), the sole North American member of its genus, is distributed in mountain forests from northern Mexico to northern British Columbia. A seed predator, *N. columbiana* is highly reliant on pine nuts from *Pinus* spp. conifers, which it both consumes directly from cones and caches for future use. Because cached seeds are often forgotten or abandoned, it is a major seed disperser for high elevation pines—in particular Whitebark Pine *P. albicaulis*. Previous studies of genetic variation in Clark’s Nutcracker found range-wide panmixia and generally high levels of heterozygosity at a handful of nuclear and mitochondrial loci, potentially due to seasonal elevational movements and long term dispersal. An earlier genome assembly from low-coverage short read data was highly fragmented and has not to date been used as the basis for population-level resequencing. Here we report on the first chromosome-scale genome assembly for *N. colombiana*. We generated long-read sequencing and genome conformation mapping data from tissues sampled from a male *N. columbiana* individual in Wyoming, USA. These data were assembled into a highly contiguous and complete assembly, which showed strong chromosomal synteny with New Caledonian crow (*Corvus moneduloides*). This genome has relatively low heterozygosity compared to other Corvid genomes and to previous population-level heterozygosity estimates for the species. We also found evidence of a long-term decline in effective population size dating back to the Pleistocene, after accounting for a technical artifact common to demographic inference using the pairwise sequential Markovian coalescent. These findings raise concerns about the future viability of the species and its mutualist *P. albicaulis*; we hope the assembly will motivate further comparative and conservation genomics research.

## Introduction

Corvids (Aves: Corvidae) are a cosmopolitan clade of passerine songbirds that includes crows and ravens, jays, and magpies. Focal and known for their intelligence and complex social and behavioral phenotypes, multiple species in the clade have served as models for research on the genomics of speciation and hybridization (Poelstra *et al*. 2014; Vijay *et al*. 2016; Knief *et al*. 2019; Slager *et al*. 2020; Metzler et al. 2021), and cognition and tool use (Sutton et al. 2018; Dussex et al. 2021). Sister to the species-rich genus *Corvus* and their close relatives jackdaws (*Coloeus* spp.), *Nucifraga* nutcrackers are a clade of four species with largely Holarctic distributions and similar seed-predator ecologies that diverged from their closest relatives at the start of the Pliocene (c.a. 5 ma) (McCullough *et al*. 2023). Prone to irruptive dispersal in search of ephemeral food resources (i.e., conifer masting events), members of the genus appear to show limited geographic differentiation (Dohms and Burg 2013, 2014; de Raad et al. 2022).

Clark’s Nutcracker (*Nucifraga columbiana*), the sole North American member of its genus, is distributed from northern Mexico to northern British Columbia, and from the Coast Range of California to the Front Range of Colorado. Highly reliant on pine nuts from *Pinus* spp. conifers (occasionally augmented with Douglas Fir *Pseudotsuga menziesii*), *N. columbiana* both consumes seeds directly and caches them for future use. Because cached seeds are often forgotten or abandoned, it is a major seed disperser for high elevation pines—particularly the five-needled Whitebark (*P. albicaulis*) and Limber (*P. flexilis*) Pines. Where the two species co-occur, the tightness of their association and Whitebark Pine’s adaptations to dispersal by *N. columbiana* has been widely described as a mutualism (Hutchins and Lanner 1982; Tomback 1982). This hypothesis and its ramifications for conservation in the face of climate change- and disease-driven declines in *P. albicaulis* have led to decades of research on diverse aspects of the behavior and evolutionary ecology of *N. columbiana* (Tomback 1982; Bednekoff and Balda 1996; Siepielski and Benkman 2007; Lorenz *et al*. 2011; Schaming 2016).

Despite well-developed genomic resources for other corvids (5 *Corvus* spp. and 1 *Aphelecoma* sp. reference genomes were available on NCBI’s Genome Data Viewer as of October 22nd, 2025; Rangwala *et al*. (2021)), *Nucifraga* nutcrackers have seen limited sequencing effort and been the focus of few population genetics studies to date. Multilocus Sanger sequencing data indicates an absence of geographic structure but substantial genetic variation (measured by observed heterozygosity *H*_*o*_ and haplotype diversity) in *N. columbiana* and what was formerly referred to as the Eurasian Nutcracker *N. caryocatactes* (Dohms and Burg 2013, 2014). A highly fragmented, short-read whole genome sequence for Clark’s Nutcracker was included as part of a broader proof-of-concept of the utility of low coverage reference genomes but included little biological interpretation of observed genetic variation, which likely included artifacts from constrained sequencing effort (Card *et al*. 2014). More recently, a study of the population genomics of *N. cayocatactes* across its range found support for three distinct, genetically isolated lineages within Eurasian Nutcracker (de Raad et al. 2022), leading to the elevation of Southern Nutcracker *N. hemispila* (itself containing four subspecies) and Kashmir Nutcracker *N. multipunctata* to species status (Gill *et al*. 2025). Despite distinct conifer mutualisms across lineages, the observed timing and pattern of divergence appeared similar to other, codistributed passerines; the authors thus downplayed local adaptation as a driver of speciation in favor of traditional Pleistocene biogeographic factors.

To aid research on corvid comparative genomics and the evolutionary ecology and conservation of *N. columbiana*, we generated a haplotype-resolved, chromosome-scale genome assembly using long-read sequencing and genome conformation mapping data from tissues sampled from a male individual collected in Wyoming, USA. Here we introduce the resource, report on technical aspects of the assembly and observed heterozygosity, and explore patterns of chromosomal synteny with New Caledonian crow *Corvus moneduloides*. We further infer demographic history from our assembly using a Pairwise Sequential Markovian Coalescent approach.

## Materials and methods

### Specimen collection and tissue sampling

We sampled heart, liver, and pectoral muscle tissues from an adult male Clark’s Nutcracker collected on October 2nd, 2023 in Medicine Bow National Forest, Albany Co., Wyoming, USA (41.239667, −105.450667). Collecting was permitted by Wyoming Game & Fish Permit (#33-754), the US Fish & Wildlife Service (Permit #MB06336A), and University of Wyoming IACUC #2022-0142. The individual was prepared as a voucher specimen (UWYMV Tissue: B-3379); it is pending accession into the University of Wyoming Museum of Vertebrates (UWYMV) ornithology collections.

### DNA extraction and sequencing

Approximately 1 gram of pectoral muscle was sent to Dovetail Genomics (Cantata Bio, LLC), where company staff isolated high molecular weight DNA using a Qiagen Genomic-tip procedure. Briefly, 100 mg of muscle tissue was first incubated with 19*µ*L RNase and 500 19 *µ*L protease in G2 lysis buffer at 50*°*C for 2 hours. 10 *µ*L of this solution was taken for total mass determination, which indicated a concentration of 1.76 *µ*g/mL. After purification and elution with a MINI column, the sample was precipitated using isopropanol, resulting in spool formation; it was then transferred to 100 *µ*L Tris-EDTA buffer and incubated overnight at 50*°*C.

The DNA sample was quantified using a Qubit 4.0 Fluorometer (Life Technologies, Carlsbad, CA, USA) and its size distribution was assessed with a Femto Pulse instrument (Agilent Technologies, Santa Clara, CA). The DNA was then sheared on a MegaRuptor 3 (Diagenode, Denville, NJ) to an average size of 15-18 kb and subsequently prepared using the SMRTbell Prep Kit 3.0 (Pacific Biosciences, Menlo Park, CA) following manufacturer’s instructions. The sequencing library was size-selected on a BluePippin instrument (Sage Science, Beverly, MA) to remove fragments smaller than 10-15 kb, bound to polymerase with the Revio polymerase kit, and sequenced in one run with a Revio sequencing plate and a Revio SMRT Cell Tray on the PacBio Revio instrument.

From the same muscle tissue, DNA for chromatin conformation capture sequencing was first fixed in place in the nucleus with formaldehyde and then extracted as described above. Fixed chromatin was digested with DNAse I, and chromatin ends were repaired and ligated to a biotinylated bridge adapter followed by proximity ligation of adapter-containing ends. After proximity ligation, crosslinks were reversed and the DNA was purified. Purified DNA was treated to remove biotin that was not internal to ligated fragments. This Dovetail Omni-C sequencing library was completed with the addition of NEBNext Ultra enzymes (New England Biolabs, Ipswich, MA) and Illumina-compatible adapters. Biotin-containing fragments were isolated using streptavidin beads before PCR enrichment of the library. It was then sequenced on an Illumina NovaSeq X Plus machine with 150 bp paired-end reads.

### Genome assembly

Prior to assembly, we removed PacBio HiFi reads with remnant adapter sequence using hifiadapterfilt (Sim *et al*. 2022), although this was probably not necessary as only 58 reads (0.00086% of total) were flagged and removed. We also estimated genome size, heterozygosity, and repetitiveness using k-mer frequency profiling of the raw HiFi data, implemented in GenomeScope2 (Ranallo-Benavidez *et al*. 2020), with the settings -l 40 -k 21 -p 2. We assembled haplotype-phased contigs with hifiasm (v0.23.0 Cheng *et al*. (2021)) using the HiFi and Omni-C reads and default settings. We ran the FCS-GX decontamination tool (Astashyn *et al*. 2024) to remove foreign contaminating sequences before scaffolding. This tool uses a custom database of NCBI RefSeq sequences and GenBank assemblies totaling *∼*700 Gbp and representing over 48,000 taxa with emphasis on prokaryotes, viruses, and eukaryotes with small genomes that are commonly found as contaminants. For hap1, 341 sequences totaling 11,798,270 bp were excluded; for hap2, 9 sequences totaling 426,226 bp were excluded. We aligned the Omni-C data to the contigs of both haplotypes and processed alignments following the Dovetail Genomics alignment pipeline (https://dovetail-analysis.readthedocs.io/en/latest/). We then used YAHS (Zhou *et al*. 2023) to scaffold the contigs.

We visualized Omni-C contact maps using PretextMap and PretextView and performed manual curation of both haplotypes following the strategy outlined previously (Howe *et al*. 2021). Overall, minimal curation was required. For hap1, we made 3 cuts in contigs, 0 breaks at gaps and 7 joins; for hap2, we made 0 cuts in contigs, 0 breaks at gaps and 6 joins.

We calculated basic assembly metrics with gfastats (Formenti *et al*. 2022). We assessed the completeness of the curated hap1 and hap2 assemblies using compleasm (Huang and Li 2023), which is a reimplementation of BUSCO (Benchmarking Universal Single-Copy Orthologs), and the passeriformes_odb12 dataset, which comprises 6,684 orthologs. We visualized chromosomal colinearity (i.e. synteny) of both assemblies compared to New Caledonian Crow reference genome (Feng *et al*. (2020); NCBI GenBank accession GCA_009650955.1) using JupiterPlot (Chu 2018), limiting the number of visualized nutcracker scaffolds to those that comprised 99% of the assembly. Reference genome GCA_009650955.1 was also used to annotate our curated hap1 assembly via liftoff (Shumate and Salzberg 2021), implemented with default settings.

We assembled a mitochondrial genome sequence with Mito-HiFi (Uliano-Silva *et al*. 2023) using the assembled scaffolds as input and a previously published Clark’s nutcracker mitogenome sequence as a reference (NCBI GenBank accession NC_022839.1; Keith Barker *et al*. (2015)). Only the hap1 assembly was found to contain mitochondrial scaffolds, which we removed to avoid redundancy with the complete mitochondrial genome. Software information and version numbers for the full assembly pipeline are provided in Table 1.

**Table 1.**
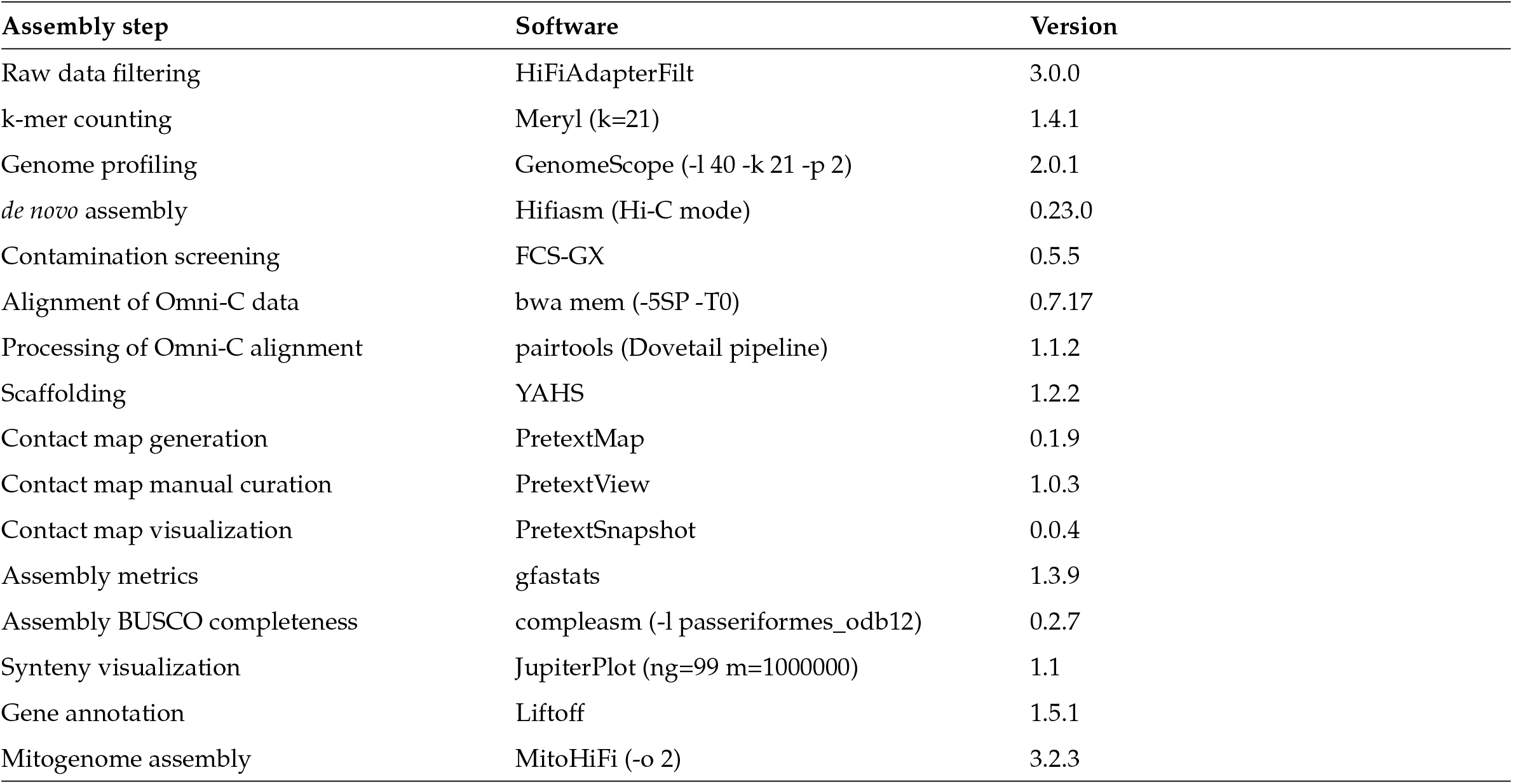
Genome assembly pipeline.

### Demographic inference

We inferred the population size history of Clark’s Nutcracker using the Pairwise Sequential Markovian Coalescent (PSMC; Li and Durbin (2011)). We aligned the PacBio HiFi reads to the completed hap1 assembly with minimap2 and used bcftools mpileup and bcftools call (Danecek *et al*. 2021) to generate a diploid consensus sequence, which was subsequently filtered for minimum and maximum read depth of 24 and 146—a third and twice the average, respectively—following PSMC recommendations. A previous study found that PSMC parameters -N30 -t5 -r5 -p “4+30*2+4+6+10” were suitable for diverse avian species (Nadachowska-Brzyska *et al*. 2015). We followed this guidance but additionally tuned the -p setting, which controls the atomic time intervals and free interval parameters. We split the first time interval in order to test for a known technical artifact identified in numerous species: a sharp population size peak immediately preceding a recent population decline (Hilgers *et al*. 2025). Specifically, we ran PSMC three times with different -p settings: (1) 4+30*2+4+6+10 (2) 2+2+30*2+4+6+10 and (3) 1+1+1+1+30*2+4+6+10. One-hundred 4 G3 Journal Template on Overleaf bootstrap replicates were performed for each analysis. Mutation rate and generation time were set to 0.3 × 10^−8^ (Vijay *et al*. 2016) and 7 years following a previous nutcracker genomic study (de Raad et al. 2022).

## Results and discussion

Our sequencing efforts produced 88.6 Gb of HiFi sequence data and 65.9 Gb of Omni-C sequence data. K-mer profiling of the HiFi data showed an estimated genome size of 1.12 Gb, with a heterozygosity of 0.38% and 15% repetitive sequence content (Fig. 1a).

**Figure 1.**
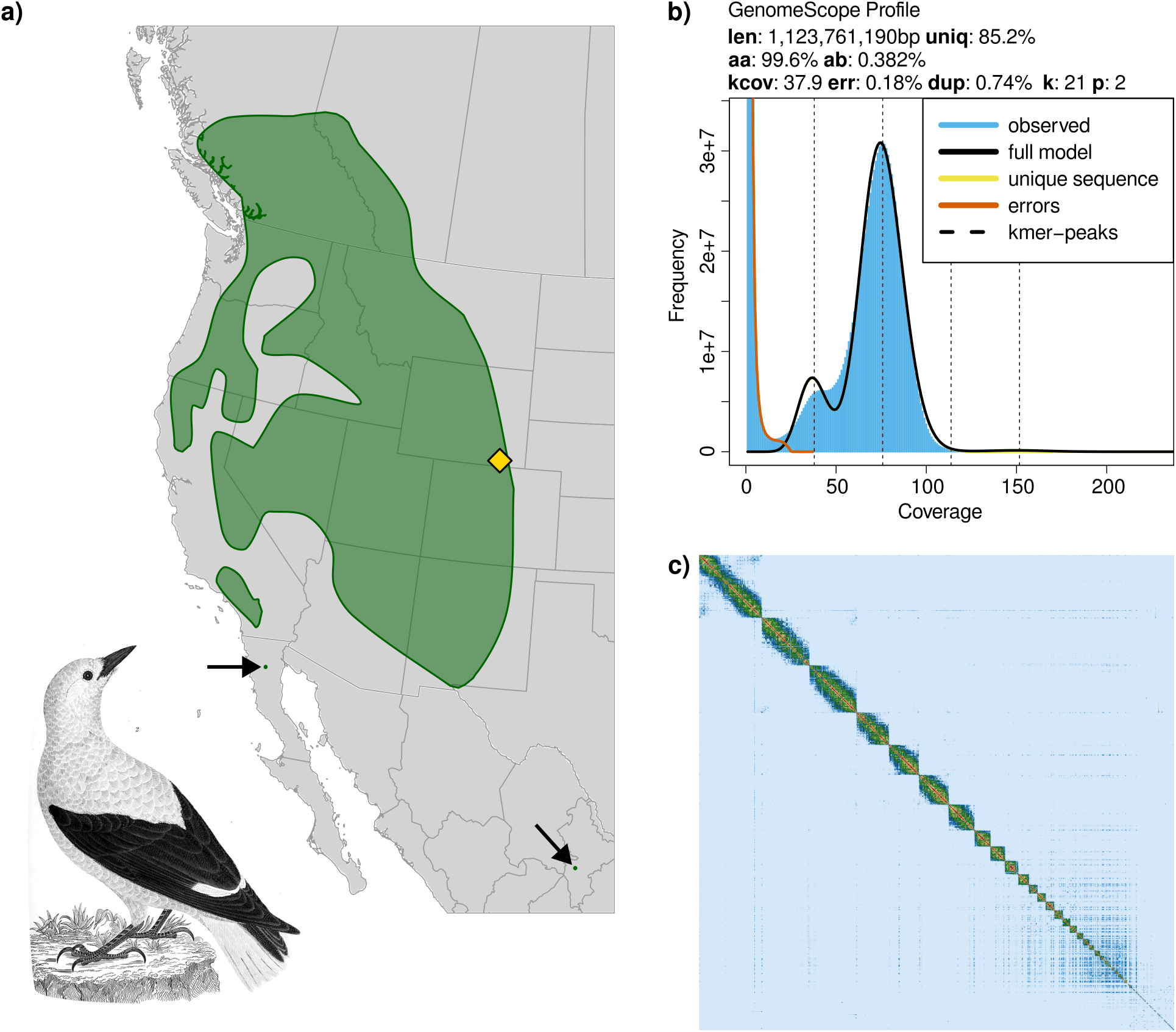
**a)** Map of the approximate year-round distribution of Clark’s Nutcracker. The sequenced bird was collected at the location marked with a yellow diamond. Distribution data was obtained from BirdLife DataZone. Illustration is by Alexander Wilson and is in the public domain in the United States. **b)** k-mer genome profiling using PacBio HiFi reads and GenomeScope2. The main peak captures average k-mer coverage of homozygous base pairs while the shorter peak (kcov = 37.9) represents that of heterozygous base pairs. Other estimated metrics are: total estimated genome length (len), percentage of genome containing non-repetitive sequence (uniq), overall rate of homozygosity (aa), overall rate of heterozygosity (ab), sequencing error rate (err), and average rate of read duplication (dup); “k” and “p” refer to settings for k-mer length and ploidy. **c)** Omni-C contact map for hap1 depicting a high-quality chromosome-scale assembly. The diagonal line represents the linear sequence of the assembly, and the image is a symmetric matrix. Frequency of contact among genomic regions, which corresponds to spatial proximity, is represented by signal intensity ranging from low (blue) to high (yellow and red).

With these data we assembled a chromosome-scale and haplotype-resolved genome sequence for Clark’s Nutcracker. Haplotype 1 comprises 806 scaffolds totaling 1.215 Gb with a contig N50 of 13.01 Mb, and haplotype 2 has similarly high contiguity (Table 2). A contact map for hap1, representing the chromosome-scale quality of the assembly, is visualized in Fig. 1b. Both assemblies contained complete and single-copy orthologs for 99.5% of the 6,684 Passeriformes BUSCO markers, suggesting highly complete assemblies (Table 2). Homology-based genome annotation relying on a closely related Corvid species, the New Caledonian Crow, resulted in 20,306 of 20,680 gene features (98.2%) successfully mapped. These contiguity and completeness metrics are among the highest for existing chromosome-scale genome assemblies in the Corvidae (Benham et al. 2023). The mitochondrial genome assembly totaled 16,896 bp, which is similar to existing *N. columbiana* and Eurasian Nutcracker *N. caryocatactes* mtDNA assemblies (16,905 and 16,914 bp, respectively; Keith Barker *et al*. (2015); Meng *et al*. (2020)).

**Table 2.**
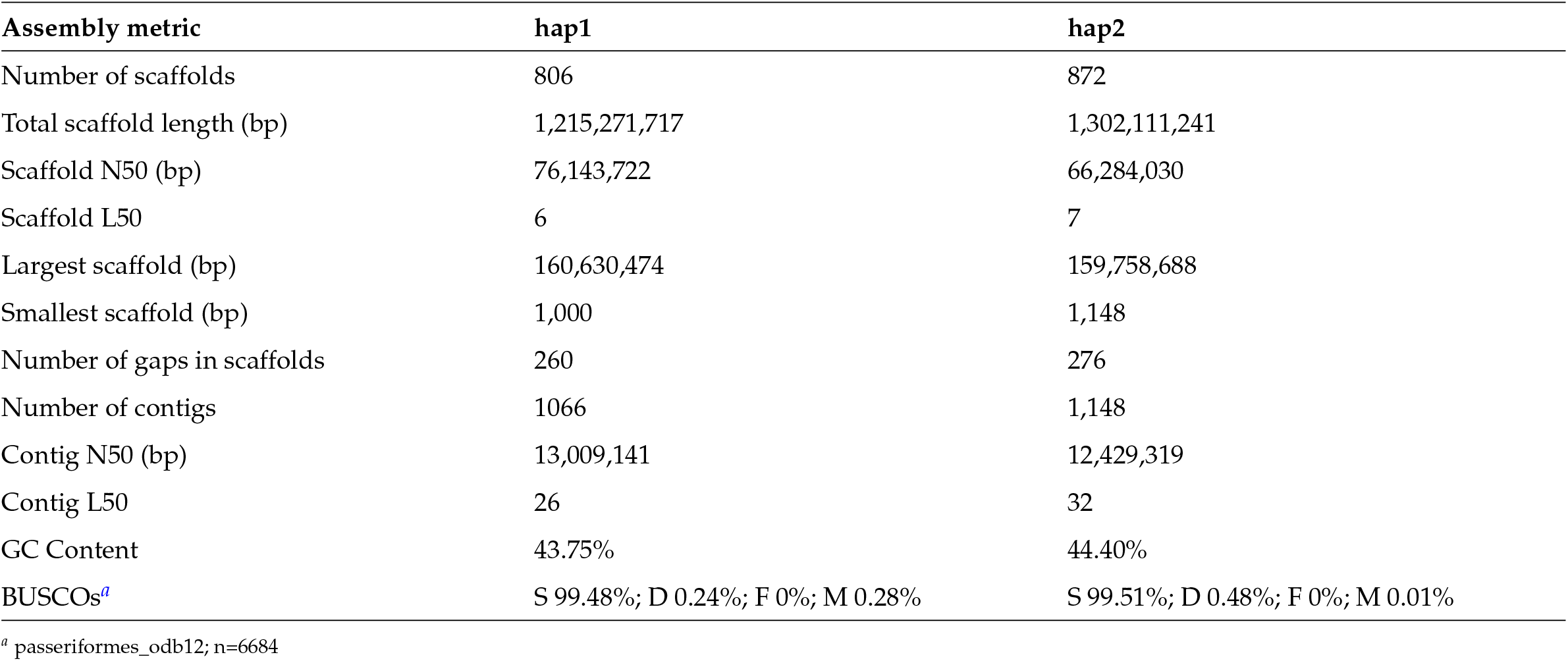
Genome assembly quality metrics.

| Assembly metric | hap1 | hap2 |
| --- | --- | --- |
| Number of scaffolds | 806 | 872 |
| Total scaffold length (bp) | 1,215,271,717 | 1,302,111,241 |
| Scaffold N50 (bp) | 76,143,722 | 66,284,030 |
| Scaffold L50 | 6 | 7 |
| Largest scaffold (bp) | 160,630,474 | 159,758,688 |
| Smallest scaffold (bp) | 1,000 | 1,148 |
| Number of gaps in scaffolds | 260 | 276 |
| Number of contigs | 1066 | 1,148 |
| Contig N50 (bp) | 13,009,141 | 12,429,319 |
| Contig L50 | 26 | 32 |
| GC Content | 43.75% | 44.40% |
| BUSCOs<sup>a</sup> | S 99.48%; D 0.24%; F 0%; M 0.28% | S 99.51%; D 0.48%; F 0%; M 0.01% |

<sup>a</sup> passeriformes\_odb12; n=6684

As expected for closely related bird species, chromosomal synteny between Clark’s Nutcracker and the New Caledonian Crow was generally high and lacked large structural rearrangements (Fig. 2). The largest gap in collinearity between assemblies occurred at the New Caledonian Crow’s W sex chromosome, corroborating notes from specimen preparation that our sequenced Clark’s Nutcracker was male, the homogametic sex (ZZ).

**Figure 2.**
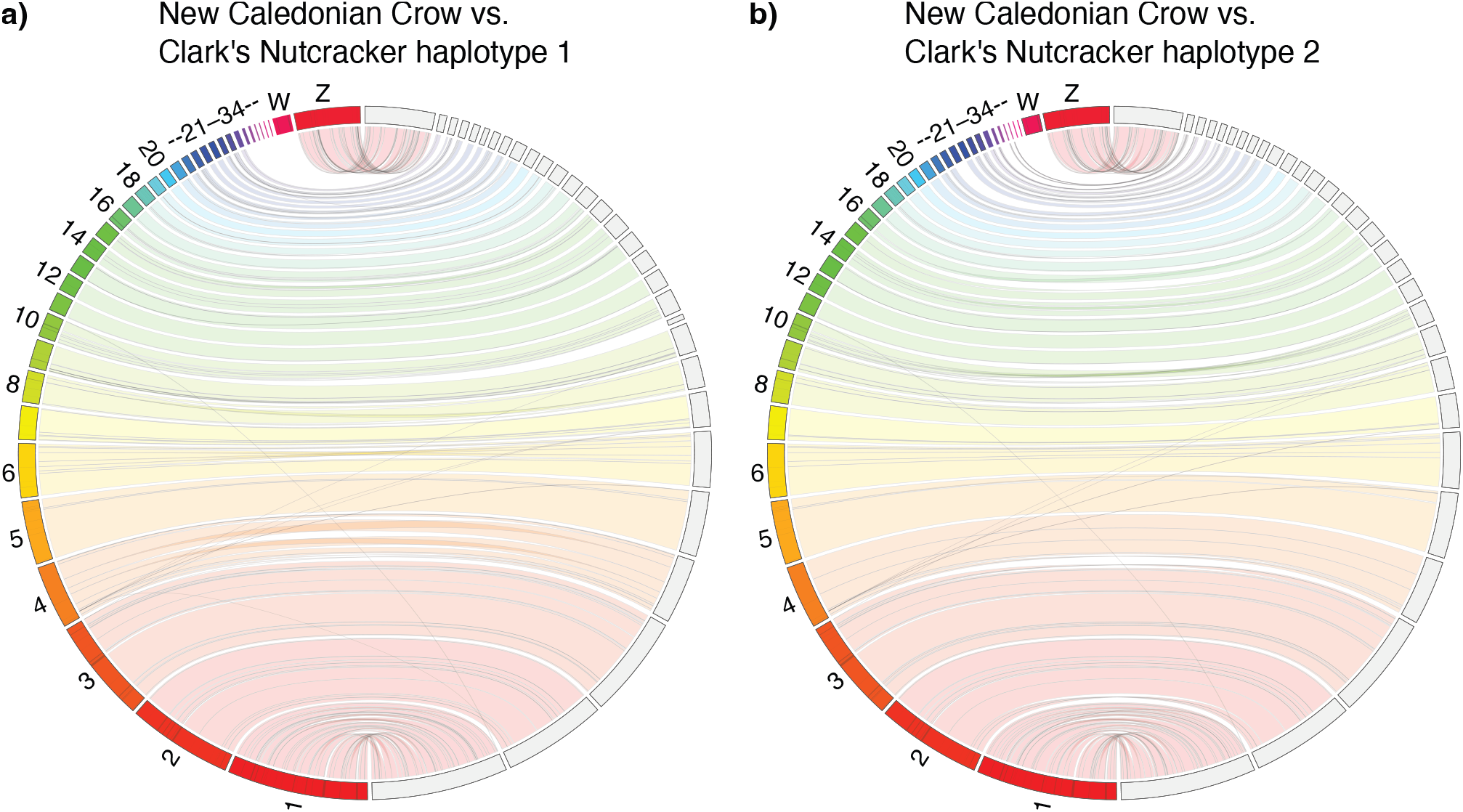
Chromosomal synteny between New Caledonian Crow and Clark’s Nutcracker, visualized with JupiterPlot. In both panels, New Caledonian Crow chromosomes are on the left hemisphere and Clark’s Nutcracker chromosomes are on the right. Colored bands indicate sequence collinearity between assemblies. The lack of homology with the New Caledonian Crow W chromosome is because a male bird was sequenced.

Inference of historical effective population size (*N*_*e*_) with PSMC suggested that Clark’s Nutcracker has experienced a long-term population decline beginning in the late Pleistocene, approximately 500 kya (Fig. 3a). We obtained this result after finding that population size history was sensitive to time interval parameter settings in a way that resembled a known technical artifact producing false population size peaks (Hilgers et al. 2025). Specifically, when the first atomic time window was kept at the default value of 4, we observed a large population size peak followed immediately by a recent decline (Fig. 3c). Splitting the first window from 4 to 2+2 removed this peak, with the exception of two outlier bootstrap replicates (Fig. 3b). Splitting the first window again (to 1+1+1+1) appeared to largely resolve the issue and revealed a consistent decline beginning *∼*500 kya (Fig. 3a). These results emphasize previous recommendations to test PSMC parameter settings beyond the default, particularly for the initial time window (Hilgers *et al*. 2025).

**Figure 3.**
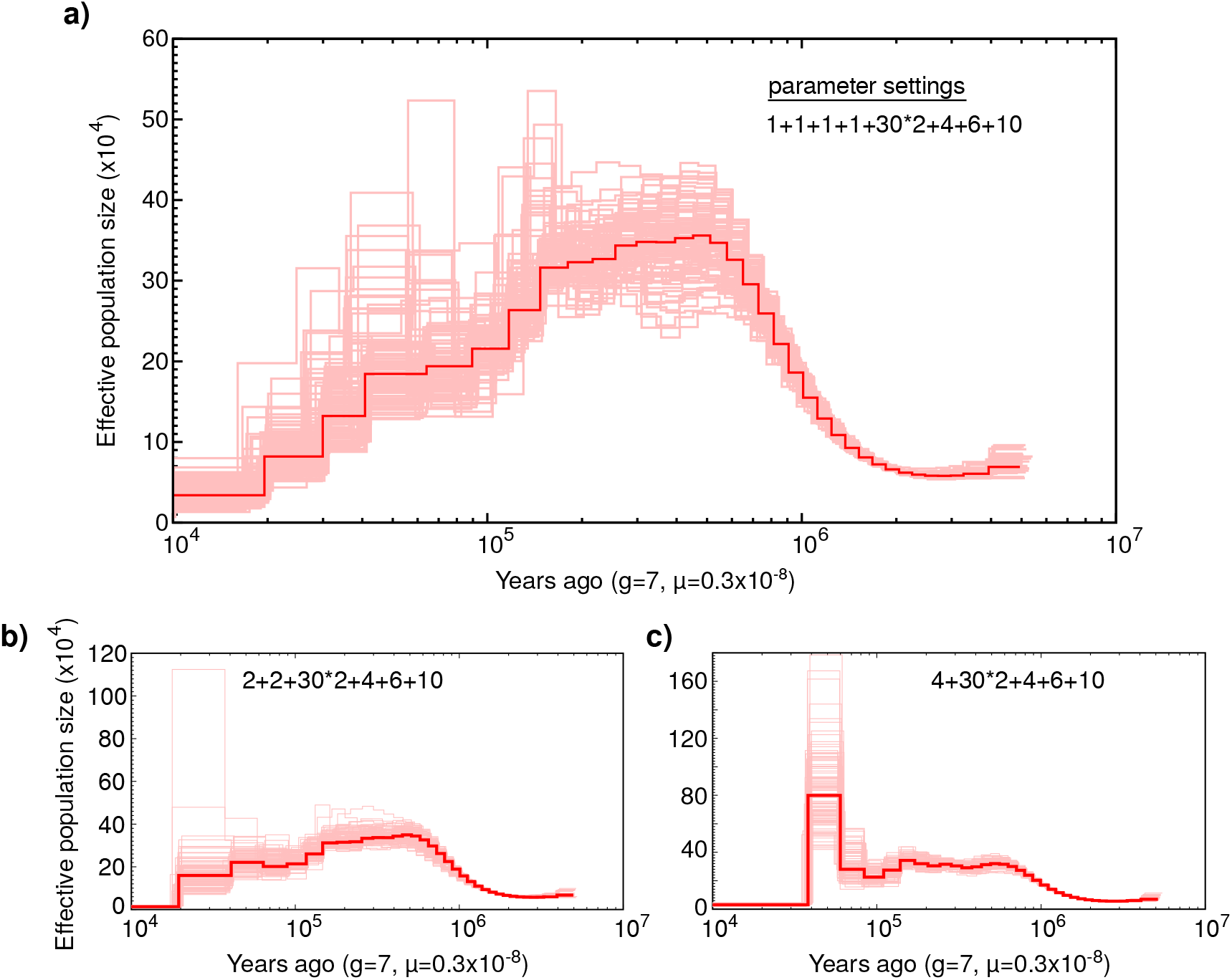
Clark’s Nutcracker population size history inferred with the Pairwise Sequential Markovian Coalescent (PSMC). The panels show the sensitivity of inference to different time interval parameter settings. **a)** The first four atomic time intervals are split into four separate windows (1+1+1+1). **b)** The first four intervals are split into two windows (2+2). **c)** The first four intervals are contained in a single window as in PSMC default settings, resulting in a false peak followed by a sharp decline.

Earlier work on the demographic history of Eurasian nutcrackers inferred recent *N*_*e*_ peaks followed by steep declines were inferred for two of three (*N. caryocatactes*) subspecies (*N. c. macrorhynchos* and *N. c. japonica*; de Raad *et al*. (2022)). We raise the possibility that these peaks might be technical artifacts, as they both closely resemble the false-peak pattern demonstrated here and by Hilgers *et al*. (2025), and as the first time interval setting in the study was kept to the problematic default value of 4. In contrast, the third Eurasian Nutcracker subspecies included in the authors’ analysis— *N. c. caryocatactes*—showed an effective population size history comparable to what we infer for Clark’s Nutcracker (Fig. 3a). Similar long-term declines from a peak in the mid-Pleistocene appear in other bird species of the temperate Northern Hemisphere (e.g. Bald Eagle, Rock Dove, and Turkey Vulture; Nadachowska-Brzyska *et al*. (2015)), and can likely be explained by dramatic shifts in climate and vegetation.

Our analysis suggests that by c.a. 10,000 years ago—roughly the end of the Last Glacial Period—the effective population size of Clark’s Nutcracker had dropped below 50,000 (Fig. 3a). A previous phylogeographic study of Clark’s nutcracker using microsatellite markers found high population-level heterozygosity, which was interpreted as a sign of large census population sizes and potential panmixia (Dohms and Burg 2013). While a direct comparison of heterozygosity between that study and ours is impossible given differences in marker type and scale (an entire genome of a single bird versus seven microsatellites in 187 birds across 13 populations), our PSMC results call into question the idea that Clark’s Nutcracker harbors high genetic diversity, as does the fact that the heterozygosity of the genome described here is notably lower than that of other Corvids such as Steller’s Jay (0.64%; Benham *et al*. (2023)) and California Scrub Jay (0.66%; DeRaad *et al*. (2023)).

We suggest that the relatively low *N*_*e*_ and *H*_*o*_ in Clark’s Nutcracker may be influenced by undescribed population sub-division or isolation-by-distance in this wide-ranging species. If irruptive movements in North America are not accompanied by gene flow and mating occurs primarily within demes structured by geography (e.g., the extent of suitable habitat in the Southern Rockies), the rate of coalescence will be determined by local *N*_*e*_, not its range-wide value. In this scenario, the male individual sequenced for this study would have large stretches of its genome that are identical-by-descent from a relatively recent ancestor that lived in or near the Laramie Range, even as a pair of alleles sampled from widely separated mountain ranges might have last shared a gamete in the much more distant past and reflect high species-level genetic diversity. Nonetheless, ongoing habitat loss during the Anthropocene and mortality in Whitebark Pine in particular lead us to believe that declining census population sizes are the primary driver of genetic diversity and *N*_*e*_ in Clark’s Nutcracker. Further population-level sequencing will be needed to examine both population genetic structure and demographic trends during the Anthropocene (Mather *et al*. 2020).

## Data availability

The raw sequence data corresponding to this genome assembly will be made publicly available upon publication on the NCBI SRA under BioProject PRJNA1347870.

## Acknowledgments

For DNA extraction and sequencing services, we thank the team at Dovetail Genomics / Cantata Biosciences, particularly Jordan Zhang and Thomas Swale. We further thank Elizabeth Wommack at UWYMV for contributions to fieldwork and curation, and Jacqueline Mattos for genome assembly guidance.

## Literature cited

Astashyn A, Tvedte ES, Sweeney D, Sapojnikov V, Bouk N, Joukov V, Mozes E, Strope PK, Sylla PM, Wagner L et al. 2024. Rapid and sensitive detection of genome contamination at scale with fcs-gx. Genome Biology. 25:60.

Bednekoff PA, Balda RP. 1996. Observational spatial memory in clark’s nutcrackers and mexican jays. Animal Behaviour. 52:833–839.

Benham PM, Cicero C, DeRaad DA, McCormack JE, Wayne RK, Escalona M, Beraut E, Marimuthu MP, Nguyen O, Nachman MW et al. 2023. A highly contiguous reference genome for the steller’s jay (cyanocitta stelleri). Journal of Heredity. 114:549–560.

Card DC, Schield DR, Reyes-Velasco J, Fujita MK, Andrew AL, Oyler-McCance SJ, Fike JA, Tomback DF, Ruggiero RP, Castoe TA. 2014. Two low coverage bird genomes and a comparison of reference-guided versus de novo genome assemblies. PLoS One. 9:e106649.

Cheng H, Concepcion GT, Feng X, Zhang H, Li H. 2021. Haplotype-resolved de novo assembly using phased assembly graphs with hifiasm. Nature Methods. 18:170–175.

Chu J. 2018. Jupiter plot: A circos-based tool to visualize genome assembly consistency.

Danecek P, Bonfield JK, Liddle J, Marshall J, Ohan V, Pollard MO, Whitwham A, Keane T, McCarthy SA, Davies RM et al. 2021. Twelve years of samtools and bcftools. GigaScience. 10:giab008.

de Raad J, Päckert M, Irestedt M, Janke A, Kryukov AP, Martens J, Red’kin YA, Sun Y, Toepfer T, Schleuning M et al. 2022. Speciation and population divergence in a mutualistic seed dispersing bird. Communications Biology. 5:429.

DeRaad DA, Escalona M, Benham PM, Marimuthu MP, Sahasrabudhe RM, Nguyen O, Chumchim N, Beraut E, Fairbairn CW, Seligmann W et al. 2023. De novo assembly of a chromosome-level reference genome for the california scrub-jay, aphelocoma californica. Journal of Heredity. 114:669–680.

Dohms KM, Burg TM. 2013. Molecular markers reveal limited population genetic structure in a north american corvid, clark’s nutcracker (nucifraga columbiana). PLoS One. 8:e79621.

Dohms KM, Burg TM. 2014. Limited geographic genetic structure detected in a widespread palearctic corvid, nucifraga caryocatactes. PeerJ. 2:e371.

Dussex N, Kutschera VE, Wiberg RAW, Parker DJ, Hunt GR, Gray RD, Rutherford K, Abe H, Fleischer RC, Ritchie MG et al. 2021. A genome-wide investigation of adaptive signatures in proteincoding genes related to tool behaviour in new caledonian and hawaiian crows. Molecular Ecology. 30:973–986.

Feng S, Stiller J, Deng Y, Armstrong J, Fang Q, Reeve AH, Xie D, Chen G, Guo C, Faircloth BC et al. 2020. Dense sampling of bird diversity increases power of comparative genomics. Nature. 587:252–257.

Formenti G, Abueg L, Brajuka A, Brajuka N, Gallardo-Alba C, Giani A, Fedrigo O, Jarvis ED. 2022. Gfastats: conversion, evaluation and manipulation of genome sequences using assembly graphs. Bioinformatics. 38:4214–4216.

Gill F, Donsker D, Rasmussen P. 2025. IOC World Bird List (v15.1). https://www.worldbirdnames.org/. Eds.

Hilgers L, Liu S, Jensen A, Brown T, Cousins T, Schweiger R, Guschanski K, Hiller M. 2025. Avoidable false psmc population size peaks occur across numerous studies. Current Biology. 35:927–930.

Howe K, Chow W, Collins J, Pelan S, Pointon DL, Sims Y, Torrance J, Tracey A, Wood J. 2021. Significantly improving the quality of genome assemblies through curation. GigaScience. 10:giaa153.

Huang N, Li H. 2023. compleasm: a faster and more accurate reimplementation of busco. Bioinformatics. 39:btad595.

Hutchins HE, Lanner RM. 1982. The central role of clark’s nutcracker in the dispersal and establishment of whitebark pine. Oecologia. 55:192–201.

Keith Barker F, Oyler-McCance S, Tomback DF. 2015. Blood from a turnip: tissue origin of low-coverage shotgun sequencing libraries affects recovery of mitogenome sequences. Mitochondrial DNA. 26:384–388.

Knief U, Bossu CM, Saino N, Hansson B, Poelstra J, Vijay N, Weissensteiner M, Wolf JB. 2019. Epistatic mutations under divergent selection govern phenotypic variation in the crow hybrid zone. Nature Ecology and Evolution. 3:570–576.

Li H, Durbin R. 2011. Inference of human population history from individual whole-genome sequences. Nature. 475:493–496.

Lorenz TJ, Sullivan KA, Bakian AV, Aubry CA. 2011. Cache-site selection in clark’s nutcracker (nucifraga columbiana). The Auk. 128:237–247.

Mather N, Traves SM, Ho SY. 2020. A practical introduction to sequentially markovian coalescent methods for estimating demographic history from genomic data. Ecology and Evolution. 10:579–589.

McCullough JM, Hruska JP, Oliveros CH, Moyle RG, Andersen MJ. 2023. Ultraconserved elements support the elevation of a new avian family, eurocephalidae, the white-crowned shrikes. Ornithology. 140:ukad025.

Meng D, Zhang Z, Li Z, Si Y, Guo Q, Liu Z, Teng L. 2020. Complete mitochondrial genome of the spotted nutcracker nucifraga caryocatactes (passeriformes: Corvidae) from shan’xi province, china. Mitochondrial DNA Part B. 5:2456–2457.

Metzler D, Knief U, Peñalba JV, Wolf JB. 2021. Assortative mating and epistatic mating-trait architecture induce complex movement of the crow hybrid zone. Evolution. 75:3154–3174.

Nadachowska-Brzyska K, Li C, Smeds L, Zhang G, Ellegren H. 2015. Temporal dynamics of avian populations during pleistocene revealed by whole-genome sequences. Current Biology. 25:1375–1380.

Poelstra JW, Vijay N, Bossu CM, Lantz H, Ryll B, Müller I, Baglione V, Unneberg P, Wikelski M, Grabherr MG et al. 2014. The genomic landscape underlying phenotypic integrity in the face of gene flow in crows. Science. 344:1410–1414.

Ranallo-Benavidez TR, Jaron KS, Schatz MC. 2020. Genomescope 2.0 and smudgeplot for reference-free profiling of polyploid genomes. Nature Communications. 11:1432.

Rangwala SH, Kuznetsov A, Ananiev V, Asztalos A, Borodin E, Evgeniev V, Joukov V, Lotov V, Pannu R, Rudnev D et al. 2021. Accessing ncbi data using the ncbi sequence viewer and genome data viewer (gdv). Genome Research. 31:159–169.

Schaming TD. 2016. Clark’s nutcracker breeding season space use and foraging behavior. PLoS One. 11:e0149116.

Shumate A, Salzberg SL. 2021. Liftoff: accurate mapping of gene annotations. Bioinformatics. 37:1639–1643.

Siepielski AM, Benkman CW. 2007. Extreme environmental variation sharpens selection that drives the evolution of a mutualism. 8 G3 Journal Template on Overleaf Proceedings of the Royal Society B: Biological Sciences. 274:1799–1805.

Sim SB, Corpuz RL, Simmonds TJ, Geib SM. 2022. Hifiadapterfilt, a memory efficient read processing pipeline, prevents occurrence of adapter sequence in pacbio hifi reads and their negative impacts on genome assembly. BMC Genomics. 23:157.

Slager DL, Epperly KL, Ha RR, Rohwer S, Wood C, Van Hemert C, Klicka J. 2020. Cryptic and extensive hybridization between ancient lineages of american crows. Molecular Ecology. 29:956–969.

Sutton JT, Helmkampf M, Steiner CC, Bellinger MR, Korlach J, Hall R, Baybayan P, Muehling J, Gu J, Kingan S et al. 2018. A highquality, long-read de novo genome assembly to aid conservation of hawaii’s last remaining crow species. Genes. 9:393.

Tomback DF. 1982. Dispersal of whitebark pine seeds by clark’s nutcracker: a mutualism hypothesis. The Journal of Animal Ecology. pp. 451–467.

Uliano-Silva M, Ferreira JGR, Krasheninnikova K, Formenti G, Abueg L, Torrance J, Myers EW, Durbin R, Blaxter M et al. 2023. Mitohifi: a python pipeline for mitochondrial genome assembly from pacbio high fidelity reads. BMC Bioinformatics. 24:288.

Vijay N, Bossu CM, Poelstra JW, Weissensteiner MH, Suh A, Kryukov AP, Wolf JB. 2016. Evolution of heterogeneous genome differentiation across multiple contact zones in a crow species complex. Nature Communications. 7:13195.

Zhou C, McCarthy SA, Durbin R. 2023. Yahs: yet another hi-c scaffolding tool. Bioinformatics. 39:btac808.

